# Assessing the response to genomic selection by simulation

**DOI:** 10.1101/2022.01.17.476687

**Authors:** Harimurti Buntaran, Angela Maria Bernal-Vasquez, Andres Gordillo, Valentin Wimmer, Morten Sahr, Hans-Peter Piepho

## Abstract

The goal of any plant breeding program is to maximize genetic gain for traits of interest. In classical quantitative genetics, the genetic gain can be obtained from what is known as “Breeder’s equation”. In the past, only phenotypic data was used to compute the genetic gain. The advent of genomic prediction has opened the door to the utilization of dense markers for estimating genomic breeding values or GBV. The salient feature of genomic prediction is the possibility to carry out genomic selection with the assistance of the kinship matrix, hence, improving the prediction accuracy and accelerating the breeding cycle. However, estimates of GBV as such do not provide the full information on the number of entries to be selected as in the classical response to selection. In this paper, we use simulation, based on a fitted mixed model for genomic prediction in a multi-environmental framework, to answer two typical questions of a plant breeder: (1) How many entries need to be selected to have a defined probability of selecting the truly best entry from the population; (2) What is the probability of obtaining the truly best entries when some top-ranked entries are selected.

## Introduction

In plant breeding programs, the breeder’s equation (Lush 1942) has been central to measure response to selection, known as genetic gain. The genetic gain also describes the breeding value of a population in one cycle of selection for the trait of interest (Rutkoski 2019). Response to selection is based on the heritability and the selection differential. Hence, a trait with a high heritability and a high selection differential can be considered to have a large genetic gain. The selection differential per se is based on the number of selected entries, which mainly relies on the breeder’s eyes when the selection is solely based on the phenotype. Environmental effects and genotype × environment interactions play a role in masking the true genotype value. This is why a breeding program is never conducted in a single environment (Crossa *et al*. 2017). The unbalancedness prevailing in datasets from the multi-environmental trials (MET) brings additional complexity to the data analysis for estimating response to selection.

The advent of genomic prediction allows breeders to estimate genomic-based breeding values via dense markers (Meuwissen *et al*. 2001). The salient feature of the genomic prediction is the improvement in the accuracy of breeding value estimation by exploiting the kinship matrix. Lorenz *et al*. (2011) and Crossa *et al*. (2017) provided a comprehensive review for the implementation and benefit of genomic prediction as the basis of genomic selection in plant breeding. Although genomic prediction can accelerate the breeding cycles and provide better accuracy in breeding value estimation, the incorporation of GEI to the genomic prediction framework is still a challenge.

Genomic prediction is by now routinely used as the basis of genomic selection in many plant breeding programs worldwide. The usual approach for evaluating the predictive accuracy of genomic prediction methods is to compute the correlation between observed and predicted genomic breeding values (GBV) using cross-validation. While this method is very useful in comparing alternative methods and designs, it does not usually give the plant breeder the full picture needed to decide on the number of breeding entries to be selected. Typical questions a plant breeder has in this regard may be exemplified as follows: (i) How many entries do I need to select to have a defined certainty (probability) of selecting the truly best entry from my population? (ii) If I select the *n* top-ranked entries, what is the probability of picking the *m* ≤ *n* truly best entries from the population?

Predictive accuracy, useful as it undoubtedly is, does not give direct answers to such crucial questions. These probabilities are hard to compute analytically for various reasons, including the unbalancedness of the data and the complexities of the mixed model used for analysis. The easiest and also the most tangible way to compute them is by simulation based on the fitted model. This idea was outlined and illustrated by examples in Piepho and Möhring (2007), and it was also used, though in slightly different context, in Piepho and van Eeuwijk (2002) and Kleinknecht *et al*. (2016). However, neither of these applications involved the use of marker data for genomic prediction. Here, we illustrate the use of this method for genomic prediction in a MET framework using an example from a hybrid rye breeding program.

## Materials and Methods

### A rye example

#### Description of the dataset and underlying population structure

The phenotypic data is proprietary of a commercial hybrid rye breeding program by KWS SAAT SE established in central Europe. In the program, the seed and pollen heterotic pools are developed and tested independently. After crossing, a few generations of single plant selection and *per se* performance evaluation, test-crosses are produced between S2-entries and two testers from the opposite heterotic pool. These test-crosses are submitted to field trials. In the first year of evaluation (hereafter GCA1 trials), entries (i.e. test-crosses) are planted in several locations.

A subset of entries is selected and forwarded to a second year of field evaluations (hereafter GCA2 trials) in more locations. In this second year of field testing, test-crosses of S_2:3_-lines (for the pollen pool) or S_3:5_-lines (for the seed pool) are crossed with four testers of the opposite heterotic pool. The selected fraction from the GCA2 trials is evaluated on the field in a third year (hereafter GCA3 trials) in more locations and additional treatments for more intensive testing. The entries of the GCA3 trials are the test-crosses developed in the previous year. A sequence of GCA1 to GCA3 trials constitute a selection cycle. All trials within a location are laid out as *α* designs with two replicates. The trial network follows a sparse pattern, where subsets of entries are evaluated in series of trials in a given subset of locations but trying to cover as many locations as possible. Sparse testing is described in Jarquin *et al*. (2020). The relatedness of the entries among cycles is high, and this breeding program can be regarded as a closed system.

Figure 1 shows the selection cycle structure of the rye hybrid breeding program. In each cycle, there are three years of tests for general combining ability (GCA) denoted here as GCA1, GCA2, and GCA3.A selection is conducted each year. Thus, in each cycle, the number of entries decreases from GCA1 to GCA3. In this study, two breeding pools were used, i.e., seed pool and pollen pool. The data available for both pools cover the years 2016 to 2020. Thus, there were three complete selection cycles available, i.e., Cycle 1 (2016 – 2018), Cycle 2 (2017 – 2019), and Cycle 3 (2018 – 2020). For Cycle 4, the available dataset comprised only GCA1-2019 and GCA2-2020, and for Cycle 5, only GCA1-2020 was available.

**Figure 1.**
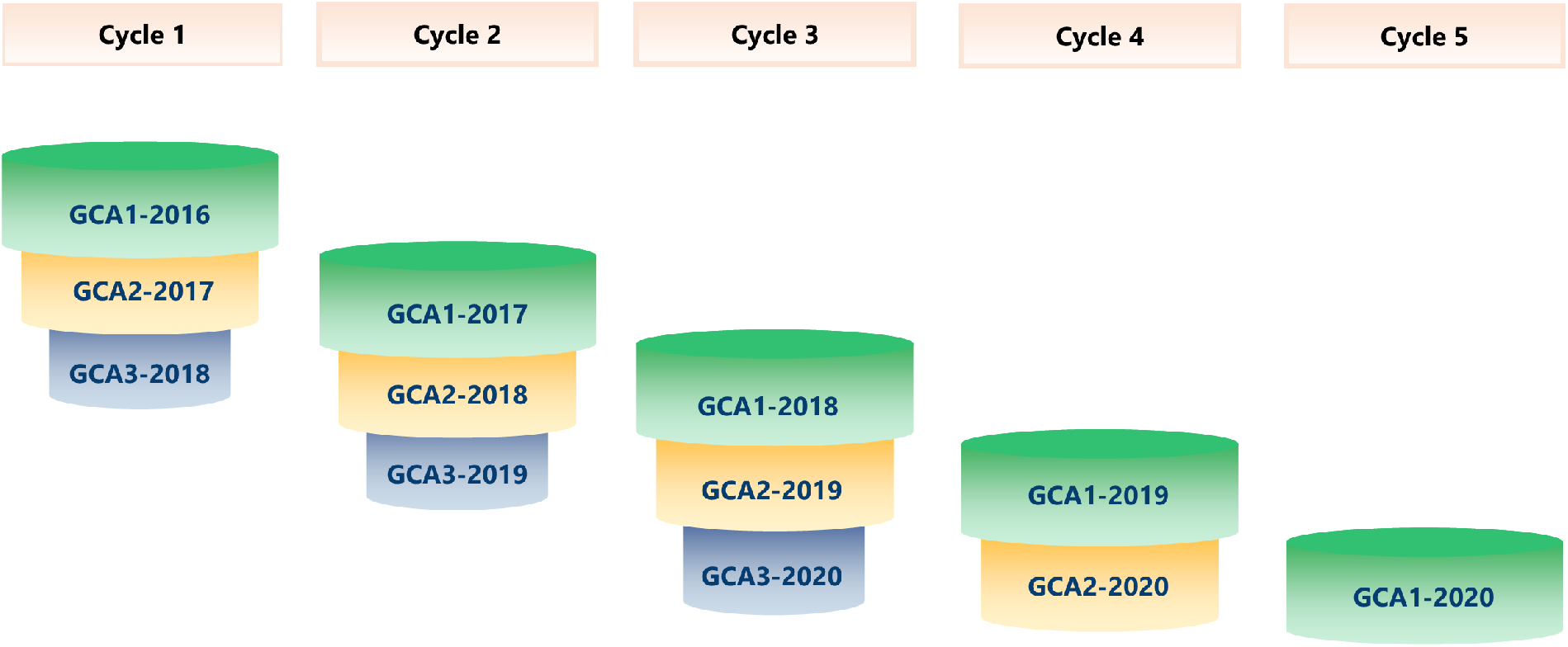
The structure of selection cycles in the rye hybrid breeding program. The number of entries decreases due to selection in each GCA trial.

#### Marker data

All genotypes from the seed and pollen pools were genotyped with an Illumina INFINIUM chip with 9963 single-nucleotide polymorphisms (SNPs) (KWS SAAT SE & Co. KG, Einbeck, Germany). The SNP used across the years partially overlap the 600 k-SNP assay of Bauer *et al*. (2017) and the 5 k-SNP assay of Martis *et al*. (2013). Monomorphic markers and markers with a minor allele frequency (MAF) <0.05, or > 10% missing values per marker were dropped. The marker cleaning was done using ASRgenomics 1.0.0 package (Gezan *et al*. 2021) implemented in R (R Core Team 2021). The number of SNP markers for the seed pool was 6246 and 7716 for the pollen pool.

### Population and statistical models

#### GCA1 response to selection assessment

The routine analysis we envision here is based on the current year’s dataset only, which in this study was taken to be the year 2020. This type of analysis is commonly done in many breeding programs, where the time between data acquisition from the current trials and selection decisions is limited. This approach also reflects the fact that selection decisions are made each year only for the entries tested in that year. We are focusing on GCA1 here, for which there is no data on the same entries from previous years.

A key challenge is that a standard single-year analysis cannot dissect the GBV×year interaction effects from GBV main effects. In particular, response to selection based on such an analysis may be over-estimated because it is based on the sum of these two effects, whereas only the first component contributes to selection response in relation to future performance of the selected entries. However, if data from multiple years is available, variance components for these two effects can be estimated, and a key idea of this paper is to use such estimates for single-year analysis in order to dissect GBV and GBV×year effects.

Estimation of the GBV×year variance from multi-year data can be done during the non-busy time of year and then plugged into the analysis for the data from the current year. In principle, we could also plug in the long-term estimate for the GBV variance estimate, but we prefer to use the current GBV variance to adjust to population structure in the current year. The major effect of the inclusion of a GBV×year interaction effect in a single-year analysis is an increased shrinkage of the genomic best linear unbiased predictions (GBLUPs) of the GBV main effects.

Moreover, the simulation based on this model will yield a smaller simulated response to selection. It is stressed here that this simulated response to selection is more realistic, as it properly reflects the fact that the apparent GBV main effect seen in a specific year based on the usual method of analysis is, in fact, the sum of the true GBV main effect and the GBV×year interaction effect for that year. In this study, the current-year (GCA1-2020) analysis uses estimates of the variances for GBV main effects and GBV×year interaction effects, which are obtained from the multi-year analysis based on the previous years. The steps for the multi-year analysis are, therefore, as follows:

1. Estimate 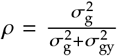 from long-term data, where 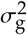 and 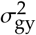 are variances of the GBV (explained in more detail in Subsection CYC and MY models) and of the GBV-by-year interaction.
2. Estimate the apparent GBV variance 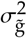 of the current year (GCA1-2020), using a model that only has a GBV main effect but no GBV×year interaction effect. It may be assumed that 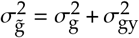.
3. Multiply this estimate of 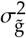 by the estimate of *ρ* to obtain an estimate of 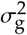 for the current year, and multiply by an estimate of (1 − *ρ*) to obtain an estimate of 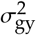 for the current year.
4. Rerun the model for GCA1-2020 data by plugging in and fixing 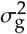 and 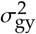 at their estimates from step 3, and using the other variance estimates obtained previously from the GCA1-2020 model. The error variance-covariance matrix of GBLUPs obtained from the rerun model will be used for the simulation, as described in more detail in Subsection The general simulation approach.

A key challenge with this approach is how *ρ* in the first step above can be estimated in the multi-year analysis. Here, we propose two possible analyses for this purpose, i.e., a combined-cycles (CYC) and a multi-year analysis based on data for GCA1 2016 – 2019 (MY). Figure 2 shows the differences in dataset structure between the CYC and MY analyses. Applying the CYC analysis, we used three complete selection cycles and one incomplete cycle, as shown in Figure 2. All entries in each GCA trial were used in the CYC analysis. On the other hand, the MY analysis only used the GCA1 datasets from the year 2016 to 2019. The checks were removed in both datasets. Table 1 summarizes the comparisons between the CYC analysis and the MY analyses regarding data handling, analysis strategy, and dataset. Moreover, the numbers of entries for the CYC and MY analyses, and the GCA1-2020 are given in Table 2.

**Table 1.**
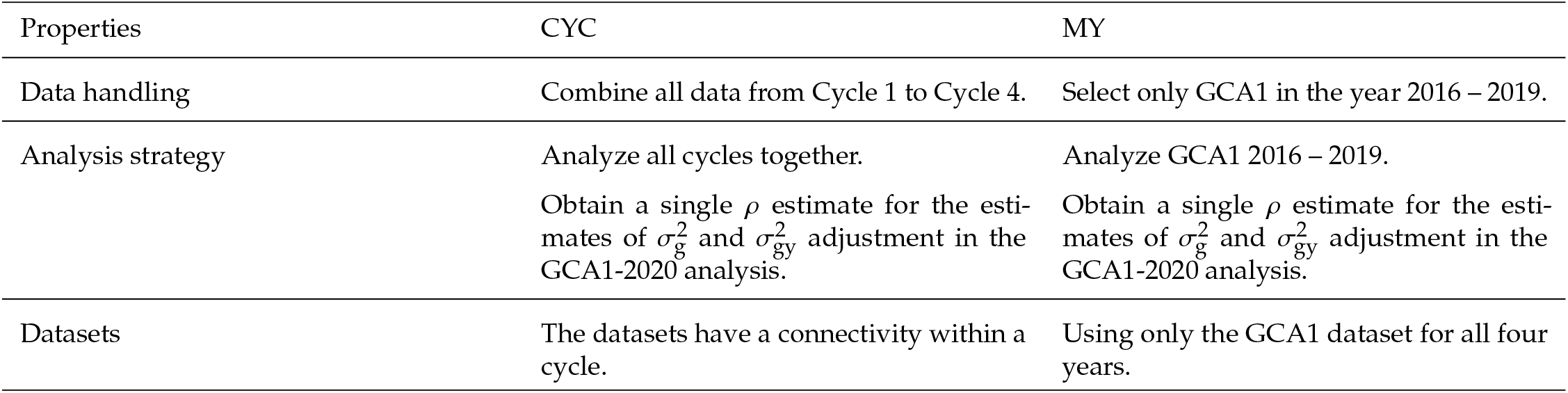
The comparisons between CYC and MY analyses in terms of data handling, analysis strategy, and datasets.

| Properties | CYC | MY |
| --- | --- | --- |
| Data handling | Combine all data from Cycle 1 to Cycle 4. | Select only GCA1 in the year 2016 – 2019. |
| Analysis strategy | Analyze all cycles together.<br>Obtain a single $\rho$ estimate for the estimates of $\sigma_g^2$ and $\sigma_{gy}^2$ adjustment in the GCA1-2020 analysis. | Analyze GCA1 2016 – 2019.<br>Obtain a single $\rho$ estimate for the estimates of $\sigma_g^2$ and $\sigma_{gy}^2$ adjustment in the GCA1-2020 analysis. |
| Datasets | The datasets have a connectivity within a cycle. | Using only the GCA1 dataset for all four years. |

**Table 2.** The number of entry levels for two analyses in the GCA1 assessment and the GCA1-2020.

| Datasets | Number of entries |  |
| --- | --- | --- |
|  | Seed pool | Pollen pool |
| CYC and MY* | 3931 | 7474 |
| GCA1-2020 | 1128 | 1910 |

\* CYC and MY have the same entry levels, but different number of observations.

**Figure 2.**
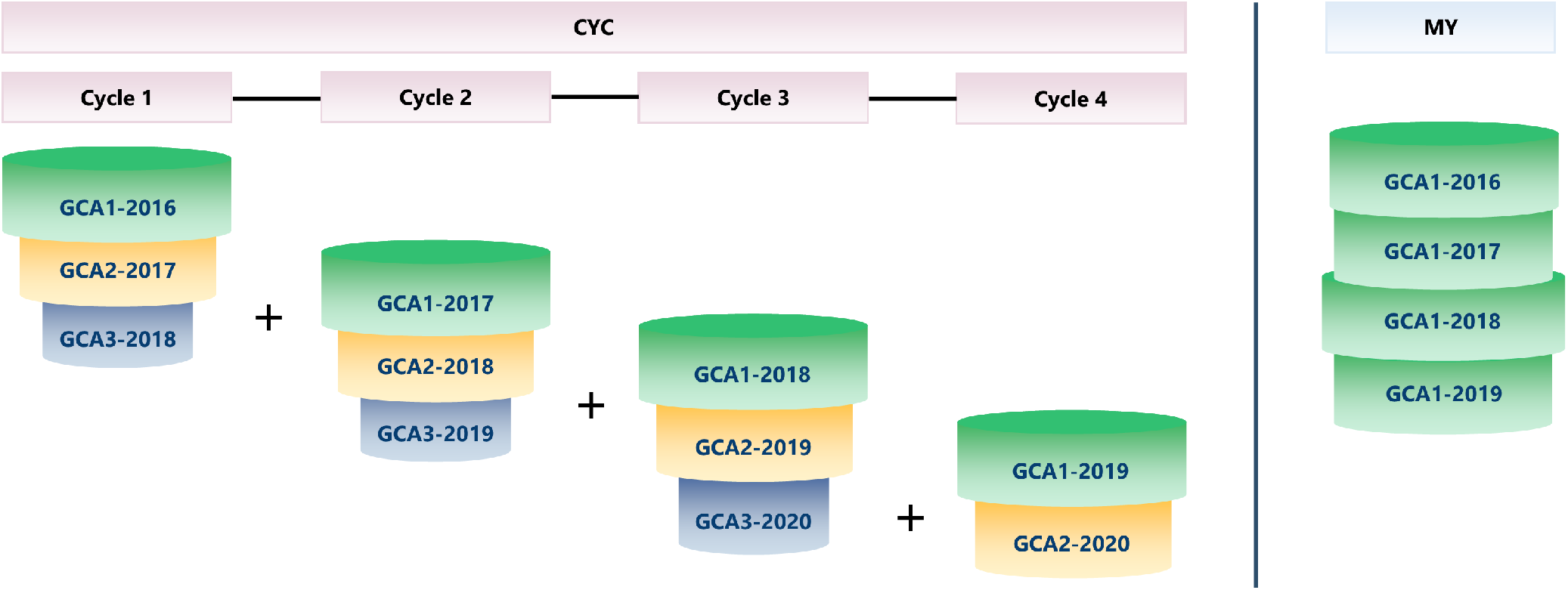
Illustration of datasets used for the CYC and MY analyses. In CYC, the dataset from all cycles are combined with each cycle comprises GCA1 – GCA3. In MY, the dataset comprises the GCA1 year 2016 to 2019. The size of the cylinders illustrates the number of entries.

### CYC and MY models

For the CYC and MY analyses, a two-stage approach was used. In the CYC analysis, the approach was applied for each cycle, while in the MY analysis, the approach was applied directly in the four-year dataset. In Stage 1, the entry means per year across locations were computed. Thus, the following linear mixed model was fitted per year for phenotypic analysis at the plot level:

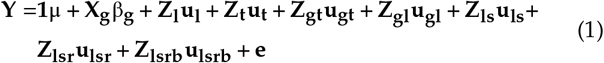

where **Y** is the vector of observed plot yields, **1** is a vector of ones, μ is the general mean, **X** is the design matrix relating to fixed effects of the entry (g), **Z** is the incidence matrix relating to random effects followed with the subscripts t, l, s, r, and b relating to the factors tester, location, trial within location, replicate within trial, and block within replicate, respectively, and **e** is the residual associated with the observation **Y**. The distributional assumption for each random effect, **u**_**l**_, **u**_**t**_, **u**_**gt**_, **u**_**gl**_, **u**_**ls**_, **u**_**lsr**_, and **u**_**lsrb**_, was a Gaussian distribution with zero mean, independence of individual effects and constant variance, as delineated in the Appendix (Table A.1). The entry effect, β_**g**_, was fixed in Stage 1 to obtain the adjusted entry means via generalized least squares, and so avoid double shrinkage (Smith *et al*. 2001; Piepho *et al*. 2012). Thus, the adjusted entry means are empirical best linear unbiased estimates (EBLUEs) of the entries’ expected values under the assumed model. The residual variance structure was heterogeneous with location-specific variance and independent error effects, as described in the Appendix (Table A.1).

In Stage 2, the adjusted entry means were assembled across several years, i.e., five years for the MY analysis and six years for the CYC analysis since it included the GCA2-2020 dataset. The following genomic prediction (GP) model was fitted at this stage:

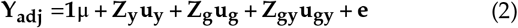

where **Y**_**adj**_ is the vector of entry-year means, **Z**_**y**_ is the incidence matrix of year main effects, **Z**_**g**_ represents the incidence matrix of entries, **u**_**g**_ is the vector of GBV, which is defined as **u**_**g**_ = **Qv**, where **Q** is the *N* × *P* marker genotypes matrix for *N* entries and *P* markers, and **v** is the vector of marker effects, 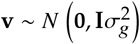, where 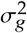 is the genomic variance. Following these assumptions, we have 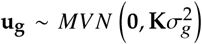 where **K** = **QQ**^**T**^. Here, **Q** was mean-centred and scaled according to the VanRaden (2008) method. There are alternative methods for computing genomic relationship matrices such as Astle and Balding (2009), Endelman and Jannink (2012), Yang *et al*. (2010), and the recent method using average semivariance by Feldmann *et al*. (2020), but these are all equivalent in terms of the resulting BLUPs of **u**_**g**_. The key component of Equation 2 is a GBV×year interaction effect, **u**_**gy**_. Thus, **Z**_**gy**_ is a block-diagonal matrix with blocks given by the coefficient of entries in a given year, 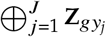, where *j* is a subscript for years, and **u**_**gy**_ ∼ *MVN* (**0, G**_gy_) is the vector of GBV-by-year effects, where 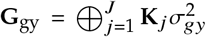, **K**_*j*_ is the kinship of all entries tested in the *j*-th year, as shown in the Appendix (Table A.2). The vector **e** is the residual term, where var(**e**) is approximated using a diagonal variance-covariance matrix as proposed in Smith *et al*. (2001). The two-stage weighted approach is useful when the single-stage approach burdens the computing time. Furthermore, a cross-validation study by Buntaran *et al*. (2020) demonstrated that the two-stage approach with Smith’s weighting was competitive to the single-stage approach.

### Current-year model

A single-stage GP model was used to obtain the apparent (unadjusted) GBV variance,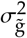. The implemented model is Equation 1 used in Stage 1 of the two-stage approach in the CYC and MY analyses, but the entry effect was replaced with the GBV as follows:

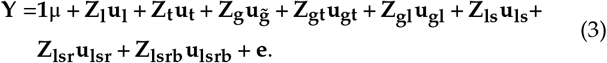

All terms are defined as for Equation 1 and explained in the Appendix (Table A.1), while 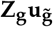 is defined as in Equation 2 and is explained in the Appendix (Table A.2). From this model, an estimate of 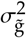 was obtained, which in its turn is used to get estimates of 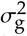 and 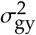 by using *ρ* from the long-term data as described in GCA1 response to selection assessment. Equation 3 was rerun with only one iteration by replacing 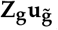 with 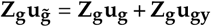. Note that the design matrix for **u**_**gy**_ is **Z**_**g**_, which is the basis of the approach suggested here. Also, in the rerun, the estimates of 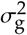 and 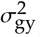 and other variance estimates of the random effects from Equation 3 were held fixed at the prespecified values.

### GCA2 response to selection assessment

The GCA2 trial is crucial since in this trial, the entries will be selected for the final trial of a selection cycle, i.e., the GCA3. Thus, it is desirable to use the GCA1 and GCA2 datasets for the GCA2 analysis. However, there is a challenge in using the previous trials’ data, i.e., the GCA1. If all entries from GCA1 are also used, then an estimate of the genetic variance applying to GCA1 is obtained (Piepho and Möhring 2006), whereas a variance estimate applying to GCA2 is needed.

Since the dropped-out entries from GCA1 have no contribution to the genetic variance in GCA2, we selected the common entries that went through to GCA2 from GCA1 as depicted with the transparent red cylinders in Figure 3. In this assessment, there were four selection cycles available from the main dataset, as shown in Figure 3. The checks were also removed as in the GCA1 assessment. The number of entries for each selection cycle is presented in Table 3.

**Table 3.** The number of entry levels for each cycle in the GCA2 assessment.

| Cycle | Number of entries |  |
| --- | --- | --- |
|  | Seed pool | Pollen pool |
| Cycle 1 | 53 | 169 |
| Cycle 2 | 81 | 160 |
| Cycle 3 | 125 | 168 |
| Cycle 4 | 94 | 172 |

**Figure 3.**
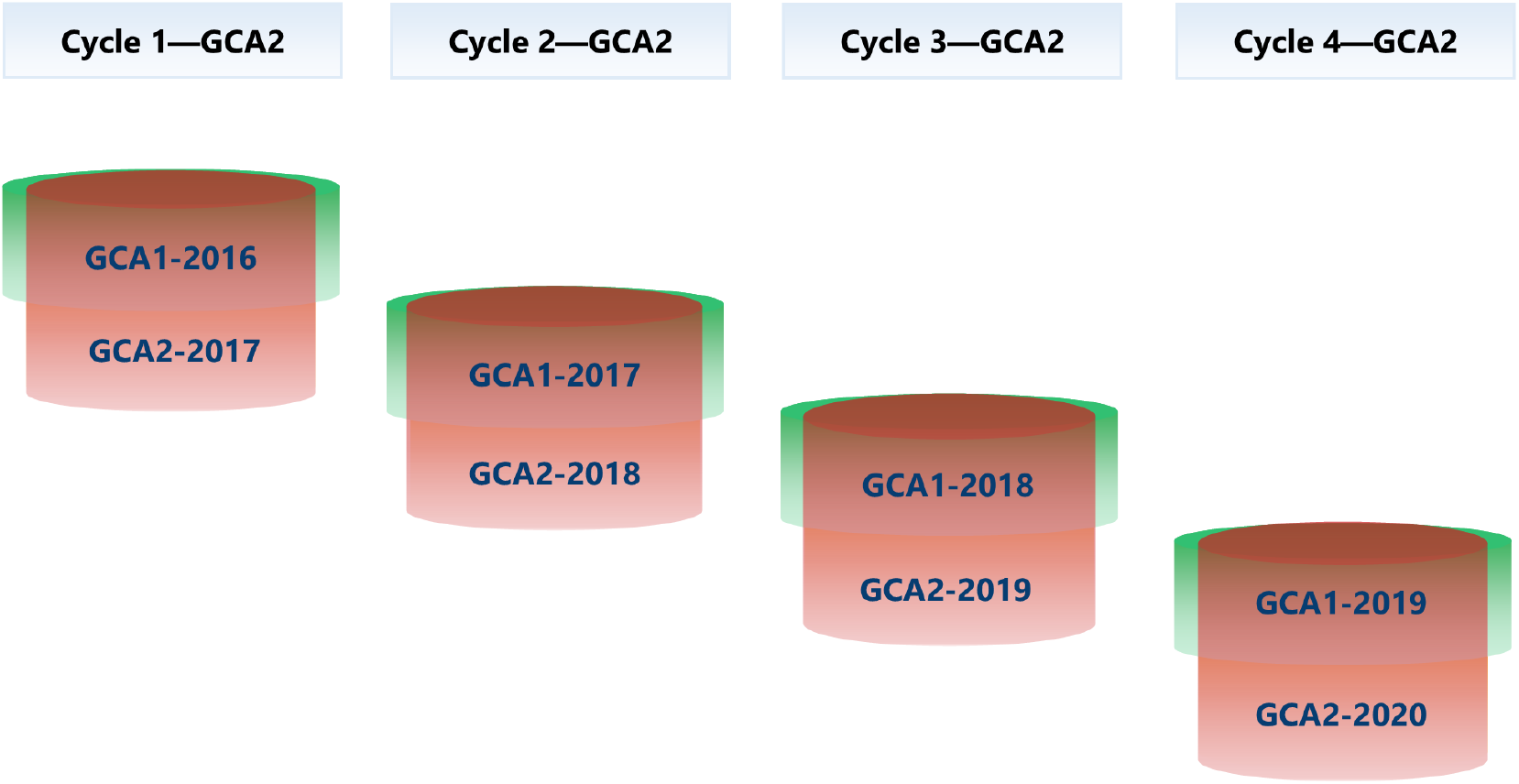
Illustration of the selected entries for the GCA2 assessment. The transparent red cylinders illustrate the selected common entries in GCA1 and GCA2.

### GCA2 model

For each cycle, a GP model was implemented using a single-stage approach. The single-stage approach was feasible to conduct due to only a relatively small number of entries in the datasets. As in the GCA1 model, the single-stage GP model for the GCA2 has a key component, i.e., a GBV×year interaction effect (**u**_**gy**_ ). The single-stage model is fitted in the plot level as follows:

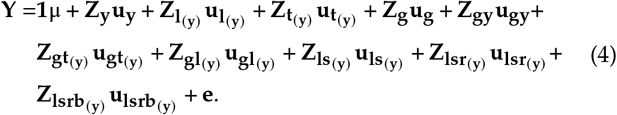

This single-stage model is used for the year-wise analysis. Thus, the tester (t), location (l), entry×tester (gt), entry×location (gl), trial within location (ls), the replication (lsr) and the incomplete block (lsrb) effects were nested within years (y). Furthermore, the distributional assumption for each random effect, 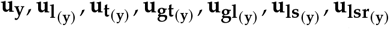, and 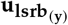 was Gaussian with zero mean, assuming independence of individual effects and year-specific variance to mimic the year-wise analysis. The GBV main effects (**u**_**g**_) and the GBV×year interaction effects (**u**_**gy**_) had the same variance-covariance structures as described in Equation 2. For the residual variance, its structure was heterogeneous with year-location-specific variance and independent error effects. The variance-covariance structure for each term in the model is explained in the Appendix (Table A.3).

### The general simulation approach

The fitted linear mixed models have a random vector **g** = (**g**_1_, **g**_2_, …, **g**_*N*_)^*T*^ of the GBV, **g**_*i*_ (*i* = 1, …, *N*) of the *N* entries in the current population from which a selection of a subset of *n* < *N* entries is to be performed. The random effects are assumed to be multivariate normal with zero mean and variance-covariance matrix 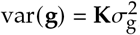, where **K** is the kinship matrix computed from markers as described in CYC and MY models, and 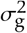 is a genomic variance. The random GBV effect is fitted within a larger linear mixed model accounting for all sources of variation due to the experimental design. In the MET framework, the model for phenotype analysis may be fitted either in a single stage or in several stages, as we demonstrate in this study. In any case, the random effects **g** will ultimately be estimated based on the GBLUPs, **ĝ**, from the fitted linear mixed models.

The key of the simulation approach is to simulate a large number *S* of realizations of the genetic effects of interest, **g**, and the corresponding estimated GBLUPs, **ĝ**, from their joint distribution and determine any quantity of interest related to the response to selection from this simulated distribution. We here use the fact that the joint distribution of **g** and **ĝ** is multivariate normal with zero mean and variance-covariance matrix

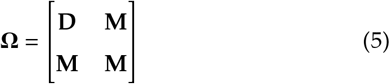

where **M** = var (**ĝ**) is the unconditional variance of (**ĝ**), 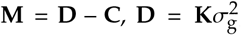, and the **C** = var(**ĝ** − **g)**, which can be obtained routinely from the inverse of the coefficient matrix of the mixed model equations (MME) (McLean *et al*. 1991; Piepho and Möhring 2007).

We use a decomposition of the **Ω** matrix given by:

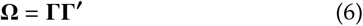

This decomposition can be obtained by Cholesky decomposition, in case the matrix **Ω** is positive-definite, or using a singular value decomposition (SVD) in the case of the **Ω** is not positive-definite.

We then simulate the values of **w**′ = (**g**′, **ĝ**′) by:

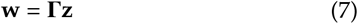

where **z** is a 2*N*-random vector of drawn from a standard normal distribution, where *N* is the size of **g**. For a single simulation of **w**, the vector **z** is generated using a random number generator, e.g., based on the Box-Muller method with the `rannor()` function in SAS, or the `rnorm()` function in R. The simulations were conducted in R 4.1.0 (R Core Team 2021) after the GCA1 and GCA2 models were fitted in ASReml-R 4.1.0.160 (Butler *et al*. 2017) using RStudio (RStudio Team 2021). The R codes for fitting the GCA1 models are provided in the electronic supplementary materials.

### Quantities of interest estimated from the simulated distribution

For each of the *S* simulation runs, the *n* best entries based on their GBLUPs **ĝ** were selected. Then, across all *S* simulation runs, we determined the proportion of cases where the selected set of *n* entries contains the *m* truly best ones based on the associated true genetic values in **g**, which is done for a range of values for *n* and *m* ≤ *n*. The probability plots showing the proportion of cases in which the selected set of *n* entries contain the *m* truly best one were generated using the `ggplot2` package Wickham (2016). Additionally, Pearson’s product-moment correlation between the **ĝ** and **g** was computed. Since the elements in **ĝ** and **g** were ranked, the correlation between the ranks of **ĝ** and the ranks of **g** was also measured. The R code for conducting the simulation and generating the probability plot is provided in the electronic supplementary materials.

## Results

### GCA1 response to selection assessment

In the GCA1 assessment, the performance of the current-year model depends on the value of 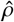 from the multi-year analysis. The estimate 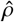 depends on the variance estimates of GBV 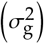 and GBV×year interaction effects 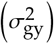. The variance estimates for year main effects 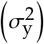, GBV main effects 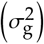, and the GBV×year interaction effects 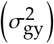 along with their associated standard errors in the CYC and MY analyses based on Equation 2 are summarised in Table 4. In the seed pool, the ratio between the estimates of 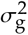 and 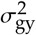 were higher in the MY analysis than in the CYC analysis because the estimate of 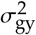 was higher in the CYC analysis than in the MY analysis. In the pollen pool, the estimate of 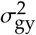 were smaller in the CYC analysis than in the MY analysis and the estimate of 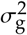 was larger in the MY analysis than in the CYC analysis. Thus, from Table 4, it can be expected that the 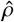 would be higher in the CYC analysis for the pollen pool and would be smaller than for the seed pool. Furthermore, in the pollen pool, the standard errors of the variance estimates of GBV did not differ much between the MY analysis and CYC analyses compared to the seed pool. On the other hand, the variance estimates of GBV×year effects were slightly higher in the MY analysis than in the CYC analysis for the pollen pool, while in the seed pool, this standard error was larger in the CYC analysis than in the MY analysis.

**Table 4.** Variance estimates and standard errors for year 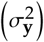, GBV 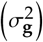, GBV×year interaction effects 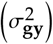 in CYC and MY based on Equation 2.

| Pool | Variance | MY |  | CYC |  |
| --- | --- | --- | --- | --- | --- |
|  |  | Estimate | SE* | Estimate | SE* |
| Seed pool |  |  |  |  |  |
| | $\sigma_y^2$ | 19.796 | 16.071 | 25.275 | 17.661 |
| | $\sigma_g^2$ | 1.960 | 0.203 | 1.279 | 0.242 |
| | $\sigma_{gy}^2$ | 1.406 | 0.151 | 4.713 | 0.279 |
| Pollen pool |  |  |  |  |  |
| | $\sigma_y^2$ | 19.882 | 16.243 | 18.02 | 12.757 |
| | $\sigma_g^2$ | 3.338 | 0.280 | 4.268 | 0.289 |
| | $\sigma_{gy}^2$ | 3.367 | 0.221 | 3.120 | 0.187 |

\* SE, standard error

The estimates and standard errors of 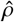 for the CYC analysis and MY analyses are given in Table 5. The standard error of the estimate of *ρ* was computed via the delta method (Lynch and Walsh 1998; Ver Hoef 2012) as implemented in the `vpredict()` function of ASReml-R (Butler *et al*. 2017). It can be seen that the differences in 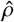 between the CYC and MY analyses were notable for the seed pool. The value of 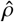 obtained from the CYC analysis (0.213) was considerably smaller than from the MY analysis (0.582), while in the pollen pool, the estimate 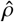 from the CYC analysis (0.578) was higher than from the CYC analysis (0.498). Moreover, the standard errors of 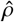 in the CYC analysis were slightly smaller than those in the MY analysis for both pools.

**Table 5.** The 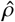 of the CYC and the MY analyses.

| Pool | MY |  | CYC |  |
| --- | --- | --- | --- | --- |
| | $\hat{\rho}$ | SE* | $\hat{\rho}$ | SE* |
| Seed pool | 0.582 | 0.044 | 0.213 | 0.038 |
| Pollen pool | 0.498 | 0.032 | 0.578 | 0.026 |

\* SE, standard error. The SE obtained via the Delta method.

The asymptotic correlations between the GBV and GBV×year variance estimates by the CYC and MY analyses for both pools given in Table 6 are negative with small values, indicating good dissection of these two variance components. The asymptotic correlation in the CYC analysis was higher than in the MY analysis in the seed pool, while in the pollen pool, the asymptotic correlation in the CYC analysis was smaller than the MY analysis. The pollen pool had smaller correlations than the seed pool, with the strongest correlation equal to -0.565. Thus, in general, there was only mild confounding between the GBV and GBV×year effects.

**Table 6.** Asymptotic correlations between the GBV and GBV×year variance estimates by the CYC and MY analyses for both pools.

| Pool | Method |  |
| --- | --- | --- |
|  | MY | CYC |
| Seed pool | -0.500 | -0.565 |
| Pollen pool | -0.469 | -0.399 |

The variance component estimates of 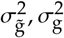, and 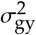 for the current year of both pools with each adjustment method are given in Table 7. The estimate of 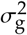 resulted in smaller values in the MY analysis for the pollen pool than with the CYC analysis, while in the seed pool, the estimates were on the opposite. Thus, the smaller 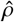 value in the CYC analysis in the seed pool led to a higher estimate of 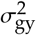.

**Table 7.**
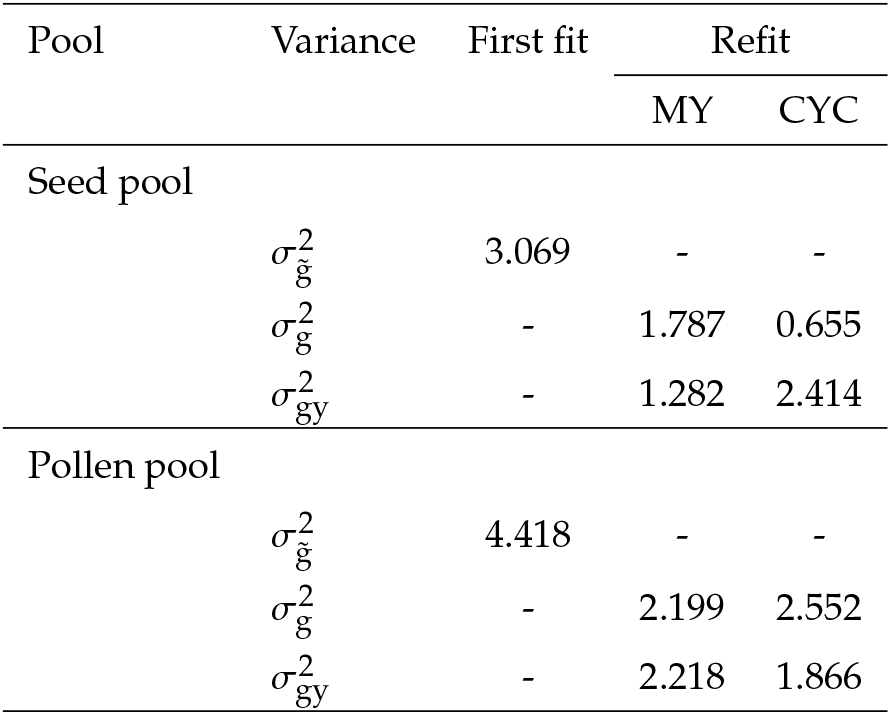
Variance estimates of the apparent GBV 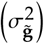, and the GBV 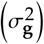, and the GBV×year interaction effects 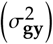 in GCA1-2020.

Table 8 presents the computing time for all analyses for GCA1. It is clear that with higher entry numbers, the computation time increased, as shown for the pollen pool. The computing time was more than 10 hours for the pollen pool on a desktop computer with an Intel i7 CPU, 64 GB RAM, and the Windows 10 (Version 21H2) 64-bit operating system. This extensive computing time was due to the presence of GBV×year interaction effect directly in the model. The computing time for the current-year analysis (GCA1-2020) was considerably longer in the pollen pool than in the seed pool due to the higher number of entries. Nevertheless, the computing time for the single-stage approach in the GCA1-2020 was still feasible. Moreover, the computing times for the simulations were trivial compared to the computing times for fitting the GP models.

**Table 8.** The computing time for Equation 2 in the GCA1 assessment on a desktop computer with an Intel i7 CPU, 64 GB RAM, and the Windows 10 (Version 21H2) operating system.

| Methods and datasets | Computing time |  |
| --- | --- | --- |
|  | Seed pool | Pollen pool |
| MY | ~2 hours | ~12 hours |
| CYC | ~2.3 hours | ~15 hours |
| GCA1-2020 (First run) | ~53 seconds | ~7 minutes |
| GCA1-2020 (Rerun and obtain $C_{22}$) | ~1.5 minutes | ~10 minutes |

The correlation coefficient between the true genetic values (g) and the GBLUPs (**ĝ)** and the correlation coefficient between *rank* (**g)** and *rank* (**ĝ)** of the GCA1 assessment for both pools based on 100K simulations are presented in Table 9. The benefit of the rank correlation is that it is not sensitive to extreme values. Also, it is concordant with the breeder’s objective to correctly rank entries. In the pollen pool, all correlation coefficients of the MY analysis were lower than the CYC analysis, while in the seed pool, all correlation coefficients of the CYC analysis were lower than the MY analysis, which was expected due to the lower ratio between the GBV variance 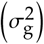, and the GBV×year 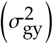 variance estimates in the seed pool’s CYC analysis as shown in Table 7. For the MY analysis, the correlation coefficients between the true genetic values **(g)** and the GBLUPs **(ĝ)**, and between *rank* **(g)** and *rank* **(ĝ)** ranged between 0.631 to 0.688 in both pools. For CYC, only in the seed pool that both correlations were relatively lower than the pollen pool. In the pollen pool, the correlations for the CYC analysis ranged between 0.682 and 0.703, while in the seed pool, the correlations ranged only 0.396 and 0.420.

**Table 9.** The correlation coefficient between the true genetic values (g) and the GBLUPs (ĝ), and the correlation coefficient between *rank* (g) and *rank* (ĝ) in the GCA1 assessment for both pools.

| Pool | Correlations | Correlation estimate |  |
| --- | --- | --- | --- |
|  |  | MY | CYC |
| Seed pool | $r(g, \hat{g})$ | 0.688 | 0.420 |
| | $r[rank(g), rank(\hat{g})]$ | 0.665 | 0.396 |
| Pollen pool | $r(g, \hat{g})$ | 0.653 | 0.703 |
| | $r[rank(g), rank(\hat{g})]$ | 0.631 | 0.682 |

The quantity of interest from the simulation is depicted as a probability plot of obtaining truly best entries for each selected proportion of the entries based on the GBLUPs, as presented in Figure 4. For a simulation with 100K iterations, the seed pool took around 1 hour and the pollen pool took around 1.5 hours on a desktop computer with an Intel i7 CPU, 64 GB RAM, and the Windows 10 (Version 21H2) operating system. The probability plots for the CYC and MY analyses are noticeably different in the seed pool, in which the probability plot for MY shows curves approaching a probability of one faster compared to the CYC analysis. The discrepancy of the probability between MY and CYC in the seed pool was due to the much smaller 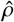 from the CYC analysis. Also, the correlation between true genetic values and GBLUPs was only 0.421, and so affected the shape of the curve for seed pool in the CYC analysis. In the CYC analysis, the probability of obtaining, the 15 truly best entries when the number of selected entries (*n*) is 50% of *N* was only around 0.21, while with the MY analysis, the probability was around 0.80. In the pollen pool, both MY and CYC showed a similar trend. Both analyses implied that by selecting a small proportion of entries, i.e., 30%, the probability of obtaining truly ten best entries was around 0.76 and 0.62 for CYC and MY analyses, respectively. Moreover, Figure 4 agrees with the correlation coefficients in Table 9, in that the higher correlation between the true genetic values and the GBLUPs, the higher the probability achieved by selecting a smaller number of entries.

**Figure 4.**
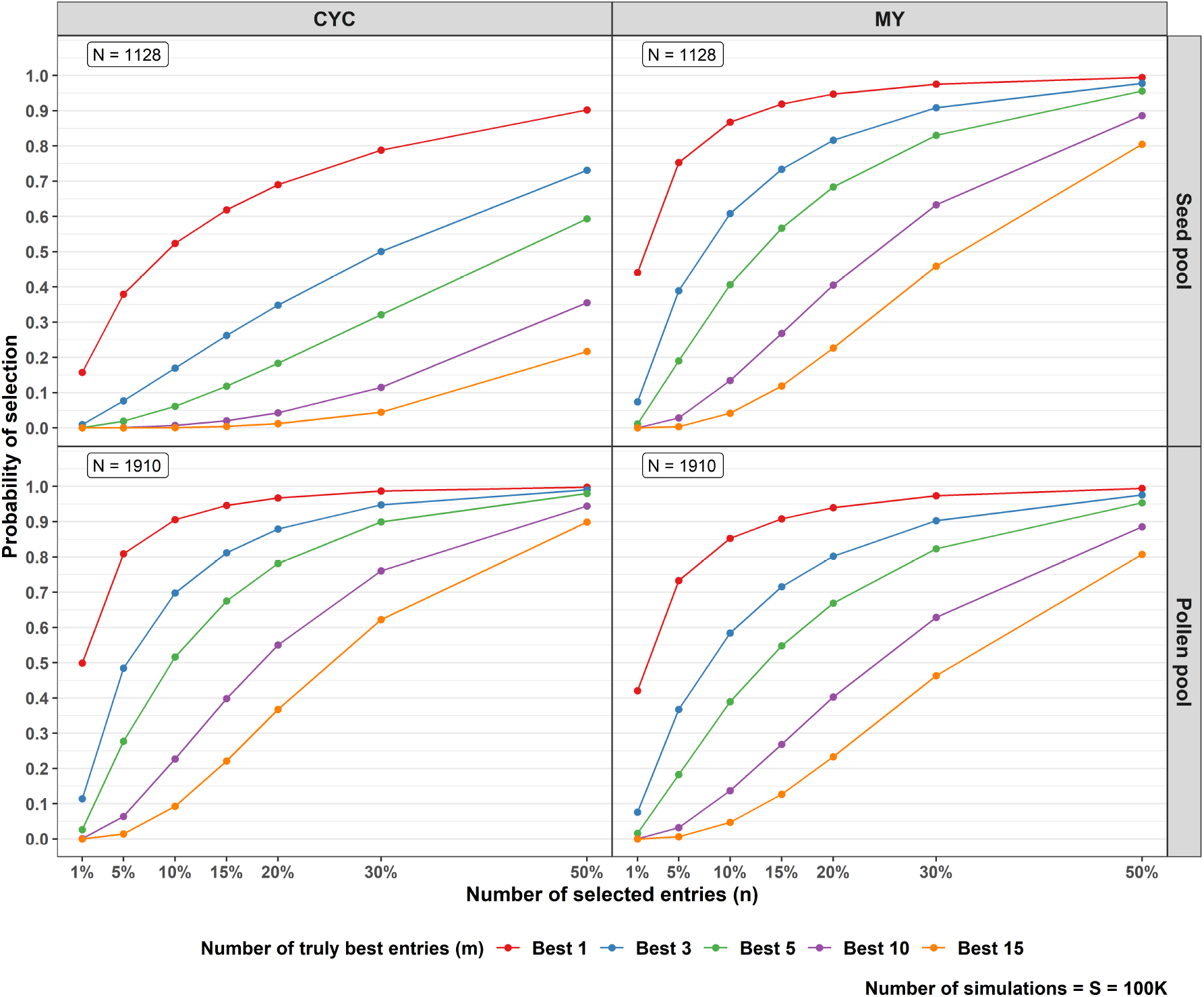
Plots of the probability of obtaining the *m* truly best entries based on the GBLUPs for each selected number of entries (expressed as percentage of *N*) from GCA1 assessment of each Pool. The different colored entries indicate the different numbers (*m*) of truly best entries.

### GCA2 response to selection assessment

The variance estimates and standard errors for year 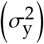, GBV 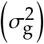, and GBV×year interaction 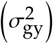 effects for each cycle are given in Table 10. In Cycle 4, the 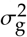 estimates were the smallest and had the smaller ratio with the 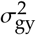 estimates for both pools. In Cycles 1 to 3, both pools had a relatively higher ratio of 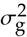 and 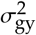 estimates.

**Table 10.** Variance estimates and standard errors for year 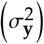, GBV 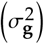, GBV×year interaction effects 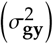 for each cycle of the GCA2 assessment.

| Pool | Variance | Estimate and standard error (SE) |  |  |  |  |  |  |  |
| --- | --- | --- | --- | --- | --- | --- | --- | --- | --- |
|  |  | Cycle 1 |  | Cycle 2 |  | Cycle 3 |  | Cycle 4 |  |
|  |  | Estimate | SE* | Estimate | SE* | Estimate | SE* | Estimate | SE* |
| Seed pool |  |  |  |  |  |  |  |  |  |
| | $\sigma_y^2$ | 9.236 | 25.546 | 6.699 | 45.889 | 0.000 | - | 0.000 | - |
| | $\sigma_g^2$ | 0.634 | 0.321 | 0.697 | 0.228 | 1.306 | 0.338 | 0.320 | 0.212 |
| | $\sigma_{gy}^2$ | 0.486 | 0.251 | 0.086 | 0.125 | 0.000 | - | 0.247 | 0.187 |
| Pollen pool |  |  |  |  |  |  |  |  |  |
| | $\sigma_y^2$ | 0.000 | - | 15.696 | 56.278 | 0.000 | - | 0.000 | - |
| | $\sigma_g^2$ | 4.269 | 0.754 | 0.707 | 0.206 | 1.707 | 0.388 | 0.396 | 0.165 |
| | $\sigma_{gy}^2$ | 0.780 | 0.305 | 0.136 | 0.111 | 0.452 | 0.200 | 0.193 | 0.141 |

\* SE, standard error.

In the seed pool, the year variance estimates in Cycle 3 to Cycle 4 were zero, while in the pollen pool, it was in the Cycle 1, Cycle 3, and Cycle 4. Furthermore, in the seed pool, the 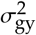 estimate in Cycle 3 was also zero. On the other hand, the year variance estimate in Cycle 2 of the pollen pool was relatively large, i.e., 15.696. Furthermore, the standard errors for the year variance estimates were large for both pools. In general, when a variance estimate goes to zero, this is possibly biased due to a small sample size. In our study, the number of years was only two years. Thus, this might be the reason that the year variance estimates were mostly zero.

The asymptotic correlations between the GBV and GBV×year variance estimates of each cycle for both pools presented in Table 11 are negative with small values. However, there was a trend that the asymptotic correlations increased from Cycle 1 to Cycle 4 in both pools. The asymptotic correlations of the seed pool were mostly higher than in the pollen pool. The highest asymptotic correlation was -0.493 in the Cycle 4 of the seed pool. In the same cycle, the asymptotic correlation for the pollen pool was -0.382. In the Cycle 3 of the seed pool, the asymptotic correlation was zero due to the variance estimate for GBV×year being zero. So, generally, there was only mild confounding between the GBV and GBV×year effects.

**Table 11.** Asymptotic correlations between the GBV and GBV×year variance estimates of each cycle of the GCA2 assessment for both pools.

| Pool | Asymptotic correlation |  |  |  |
| --- | --- | --- | --- | --- |
|  | Cycle 1 | Cycle 2 | Cycle 3 | Cycle 4 |
| Seed pool | -0.400 | -0.268 | 0.000 | -0.493 |
| Pollen pool | -0.205 | -0.307 | -0.333 | -0.382 |

Table 12 presents the computing time of each cycle analysis for each pool. Due to relatively much smaller number of entries, the seed pool took around 2 to 5 seconds, while in the pollen pool, it took around 13 to 15 seconds. Furthermore, the GCA2 analyses were done using a single-stage approach, which was feasible due to a small number of entries and the number of years being only two.

**Table 12.** The computing time for all analyses in the GCA2 assessment on a desktop computer with an Intel i7 CPU, 64 GB RAM, and the Windows 10 (Version 21H2) operating system.

| Pool | Computing time |  |  |  |
| --- | --- | --- | --- | --- |
|  | Cycle 1 | Cycle 2 | Cycle 3 | Cycle 4 |
| Seed pool | ~2 seconds | ~4 seconds | ~5 seconds | ~4 seconds |
| Pollen pool | ~13 seconds | ~13 seconds | ~15 seconds | ~15 seconds |

Table 13 provides the correlation coefficients between the true genetic values (**g)** and the GBLUPs (**ĝ)** and the correlation coefficients between *rank* (**g)** and *rank* (**ĝ)** of each cycle of the GCA2 assessment for both pools based on 100K simulations. The lowest correlation coefficients for both pools were observed in Cycle 4 due to smaller 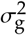 estimates and their ratio to the 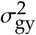 estimates compared to the other cycles. The correlation coefficient between **g** and **ĝ** was 0.616 and 0.672 for the seed and pollen pools, respectively. These coefficients were much lower than the correlation coefficients in other cycles in both pools. The pattern of the correlation coefficients of the rank of **g** and **ĝ** was the same as the correlation coefficients between **g** and **ĝ**.

**Table 13.** The correlation coefficient between the true genetic values (g) and the GBLUPs (ĝ), and the correlation coefficient between *rank* (g) and *rank* (ĝ) in the GCA2 assessment for both pools.

| Pool | Correlations | Correlation estimate |  |  |  |
| --- | --- | --- | --- | --- | --- |
|  |  | Cycle 1 | Cycle 2 | Cycle 3 | Cycle 4 |
| Seed pool | $r(g), (\hat{g})$ | 0.736 | 0.820 | 0.865 | 0.616 |
| | $r[rank(g), rank(\hat{g})]$ | 0.700 | 0.790 | 0.839 | 0.578 |
| Pollen pool | $r(g), (\hat{g})$ | 0.899 | 0.804 | 0.837 | 0.673 |
| | $r[rank(g), rank(\hat{g})]$ | 0.883 | 0.778 | 0.816 | 0.645 |

The quantity of interest from the simulation for GCA2 assessment is depicted as a probability plot of obtaining truly best entries given the number of selected entries in Figure 5. The 100K simulations took around 30 minutes for the seed pool and took around 40 minutes the pollen pool on a desktop computer with an Intel i7 CPU, 64 GB RAM, and the Windows 10 (Version 21H2) operating system. As we can see, the differing number of entries played a significant role. The pollen pool had a relatively larger number of entries compared to the seed pool. The probability of obtaining the truly best entry was, therefore, higher, especially in Cycle 1. In Cycle 1, due to a much lower number of entries in the seed pool, selecting 20% entry result in a probability of nearly 0 of having picked the ten truly best entries, while in the pollen pool, the probability was around 0.6. In Cycle 4, although the number of entries in both pools was decent, both pools had relatively low ratios of the GBV and GBV×year variance estimates as shown in Table 10, and so the correlations between true GBV and GBLUPs were relatively low, as shown in Table 13. Thus, the probability of obtaining truly best entries was relatively low compared to Cycles 2 and 3. For example, having the ten truly best entries by selecting 20% of the entries only achieved a probability of around 0.06 for the pollen pool and nearly 0 for the seed pool.

**Figure 5.**
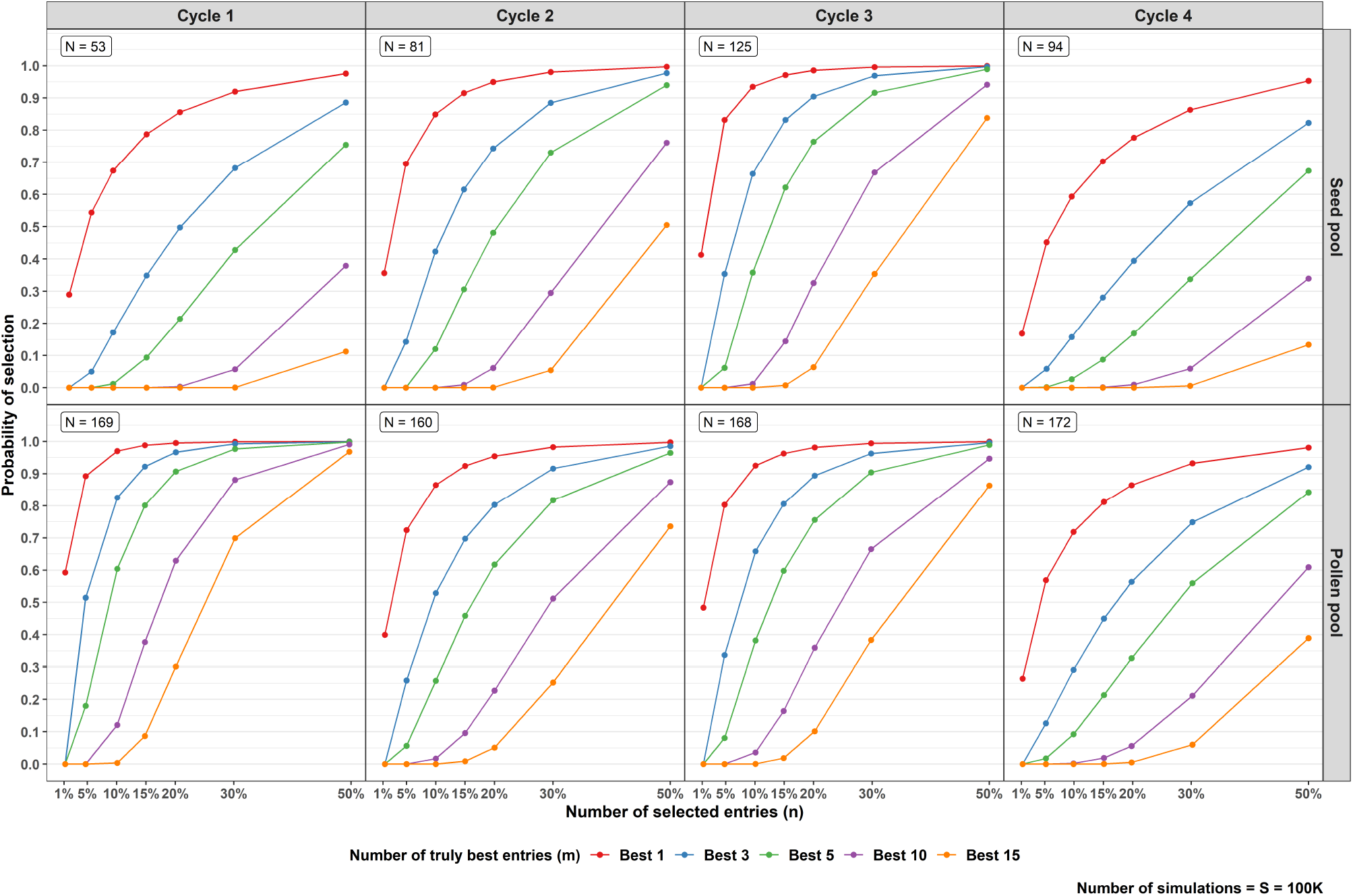
Plots of the probability of obtaining the *m* truly best entries based on the GBLUPs for each selected number of entries (expressed as percentage of *N*) of each cycle in the GCA2 assessment for each Pool. The different colored entries indicate the different numbers (*m*) of truly best entries. In general, the pollen pool has higher probability to obtain the truly best entries than the seed pool. In Cycle 4, both pools have a relatively lower probability to obtain the truly best entries compared to the other cycles.

## Discussion

The response to selection has been widely used in plant breeding programs to measure genetic gain. Our proposed simulation allows breeders to obtain information on how well the selection can be made for the next selection stage in the genomic selection framework. The response to selection can be measured in terms of the probability of selecting the truly best entry based on the number of selected entries.

In the GCA1, the number of entries is normally large since this is the first year that all the entries enter the GCA testing trial after a per-se line selection stage for agronomic (non-yield) traits. In comparison to using only GCA1 from the years 2016 to 2019 as in the MY analysis, the CYC analysis benefits from using the entries in the selected fractions of GCA2 and GCA3 improving the connectivity across GCA trials. In the same vein, Smith *et al*. (2021) also reported that the poor connectivity could lead to poor estimated genetic variance parameters, and so decreasing the genetic gain (Sales and Hill 1976a,b). Furthermore, the use of the kinship matrix to model GBV×year interactions can be advantageous, as Bernal-Vasquez *et al*. (2017) reported. By using the kinship, we gained more connectivity since we were estimating the individual marker effects. Thus, even when the entries were only available in one year, the marker allele appeared on more than one year.

Discrepancies between MY and CYC in the pollen pool were less pronounced compared to the seed pool. The connectivity in the CYC analysis slightly enhanced the estimate of *ρ* in the pollen pool but not in the seed pool, which might be due to the genetic sampling in the seed pool population. In the seed pool, the CYC analysis had higher probability of bias because, although the GCA2 and GCA3 entries improve connectivity, the selected fraction may be composed of lines that trace back in their pedigree to only very few parental components (predominant family selection) that show a very particular change of ranking between years, which leads to an over-estimation of GBV×year and an underestimation of *ρ*. Furthermore, Table 4 shows that the GBV variance estimates of the seed pool are far smaller compared to the pollen pool in both MY and CYC, which indicates that there was unbalanced recombination in the seed pool. The risk of such a bias is expected to be potentiated when the environmental conditions change considerably between years. Here, the year of 2017 was extremely wet, whereas the years of 2018, 2019, and 2020 were extremely dry. Such effects are compounded and when there is a genetic bottleneck in the breeding population.

We have also shown that the simulation for the GCA2 assessment is useful to measure the selection accuracy for the final trial in the GCA3. Thus, the simulation was only based on the selected entries that progressed to GCA2, which was the orthogonal kernel of the GCA1 and GCA2. The simulations results showed that when the 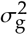 estimate was relatively larger than the 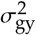 estimate, the probability of obtaining the truly best entry based on the GBLUPs was high, which shown by Cycles 2 and 3 in the pollen pool. Using only the GCA2 data will lead to a relatively small amount of data and the GBV×year interaction effect cannot be dissected. Thus, it may be still very beneficial to joint data from the different trials.

Kleinknecht *et al*. (2016) performed similar simulations using all the datasets and simulated a selection in each year, while our simulation did not simulate the selection in each year. The rationale was that the actual selection is a complex decision, i.e., based on multiple traits, criteria and considerations, often not encoded in a way that can be simulated but mostly resulting from a final integrated breeder’s-eye decision.

In this study, the number of years was not large, i.e., only 4 years in the GCA1 assessment, and the simulations were conducted using the estimates produced by frequentist residual maximum likelihood method (REML). Furthermore, as demonstrated by our results, estimation of the variance components for GBV and GBV×year effects is the Achilles’ Heel of the whole approach. These variance components are expected to display year-to-year variation, which is why our approach prescribes re-estimation of each variance in a new year using a long-term estimate of *ρ*. The expected year-to-year variability, however, suggests that a fully Bayesian approach that operates on a prior distribution for each variance component, rather than a fixed value, may be beneficial. The key challenge with such a framework is how to properly inform these priors and the need to integrate long-term data from an ongoing breeding program.

An approach that can be beneficial is to continuously use a Bayesian framework to collect more information from the previous years and use it as a prior information to update the current-year analysis. This approach is known as Bayesian Updating (Sorensen and Gianola 2002). In Bayesian Updating, the prior distribution is based on the previous posterior distribution. In this case, the Bayes theorem has “memory”, and the inferences can be updated sequentially. For example, for the GCA1 analysis, the prior distribution for the current-year analysis can be obtained from the posterior distribution of many previous GCA1 or Cycle analyses. Therefore, a further study with the Bayesian Updating framework would be worthwhile to investigate in the future.

Furthermore, the Gibbs sampling for estimating variance components might be appealing compared to REML since it used prior distribution that could produce more accurate variance estimates (Van Tassell et al. 1995). In the GCA2 assessment, we only chose the full set entries that progressed to GCA2 from GCA1, and so we did not face the missing-not-at-random pattern that can lead to a bias by using REML (Piepho and Möhring 2006; Hartung and Piepho 2021). However, the year variance estimates were mostly zero in the GCA2 assessment, resulting from a non-negativity constraint on REML estimates. In this study, the zero estimates of the year variance might be due to a small number of years. Modelling the year effect as fixed can be an alternative, as shown in Table S.1 in the Supplementary Tables. In Table S.1, the GBV and GBV×year variance estimates are very close to their estimates in Table 4. Thus, the estimates of *ρ* in Table S.2 were also very similar to the estimates in Table 5. The similar pattern also showed for the GCA2 assessment. The variance component estimates of GBV and GBV×year in Table 10 and in Table S.3 of the Supplementary Tables are very similar. Thus, modelling the year effect as fixed can be an alternative option.

Despite issues around the best dataset for estimating the variance components, we demonstrated that the genetic gain can be measured through simulations in genomic prediction framework, either using CYC or MY analyses. We recommend to choose the analysis based on the breeding population structure. The MY analysis is suitable when the population structure is highly influenced by strong selection intensity, e.g., the seed pool, while the CYC analysis can be chosen when there is no pre-dominant selection, e.g., the pollen pool. Further studies to compare the MY and CYC approaches using other crops and breeding populations is, therefore, worthwhile to be conducted.

## Appendix

**Table A.1.** The variance–covariance structures for each term in Equation 1 and Equation 3 for CYC strategy.

| Term and assumption | Variance-covariance structure * | Remarks |
| --- | --- | --- |
| $\mathbf{u}_l \sim N(\mathbf{0}, \mathbf{G}_l)$ | $\mathbf{G}_l = \mathbf{I}\sigma_l^2$ | Identity variance structure for each random effect factor. |
| $\mathbf{u}_t \sim N(\mathbf{0}, \mathbf{G}_t)$ | $\mathbf{G}_t = \mathbf{I}\sigma_t^2$ | |
| $\mathbf{u}_{gt} \sim N(\mathbf{0}, \mathbf{G}_{gt})$ | $\mathbf{G}_{gt} = \mathbf{I}\sigma_{gt}^2$ | |
| $\mathbf{u}_{gl} \sim N(\mathbf{0}, \mathbf{G}_{gl})$ | $\mathbf{G}_{gl} = \mathbf{I}\sigma_{gl}^2$ | |
| $\mathbf{u}_{ls} \sim N(\mathbf{0}, \mathbf{G}_{ls})$ | $\mathbf{G}_{ls} = \mathbf{I}\sigma_{ls}^2$ | |
| $\mathbf{u}_{lsr} \sim N(\mathbf{0}, \mathbf{G}_{lsr})$ | $\mathbf{G}_{lsr} = \mathbf{I}\sigma_{lsr}^2$ | |
| $\mathbf{u}_{lsrb} \sim N(\mathbf{0}, \mathbf{G}_{lsrb})$ | $\mathbf{G}_{lsrb} = \mathbf{I}\sigma_{lsrb}^2$ | |
| $\mathbf{e} \sim N(\mathbf{0}, \mathbf{R})$ | $\mathbf{R} = \bigoplus_{l=1}^L \mathbf{R}_l$<br>$\mathbf{R}_l = \mathbf{I}\sigma_{\varepsilon_l}^2$ | Heterogeneous location-specific consisting of a block-diagonal matrix with diagonal element $\sigma_{\varepsilon_l}^2$, $l = 1, 2, \dots, L$. |

\* The subscript $l$ denotes the $l$ -th location; $L$ is the number of location.

**Table A.2.** The variance–covariance structures for each term in Equation 2 and Equation 3 for CYC strategy.

| Term and assumption | Variance-covariance structure * | Remarks |
| --- | --- | --- |
| $\mathbf{u}_g \sim MVN(\mathbf{0}, \mathbf{K}\sigma_g^2)$ | $\mathbf{K}\sigma_g^2$ | $\mathbf{u}_g = \mathbf{Q}\mathbf{v}$, where $\mathbf{Q}$ is the marker genotypes matrix, $\mathbf{v}$ is the vector of marker effects, $\mathbf{v} \sim N(\mathbf{0}, \mathbf{I}\sigma_g^2)$, where $\sigma_g^2$ is the genomic variance.<br>Based on these assumptions $\mathbf{u}_g \sim MVN(\mathbf{0}, \mathbf{K}\sigma_g^2)$, where $\mathbf{K} = \mathbf{Q}\mathbf{Q}^T$.<br>The $\mathbf{K}$ matrix is the genomic relationship. |
| $\mathbf{u}_{gy} \sim MVN(\mathbf{0}, \mathbf{G}_{gy})^\dagger$ | $\mathbf{G}_{gy} = \bigoplus_{j=1}^J \mathbf{K}_j\sigma_{gy}^2$ | The matrix $\mathbf{Z}_{gy}$ is a block-diagonal matrix with blocks given by the coefficient of entries in a given year, $\bigoplus_{j=1}^J \mathbf{Z}_{gy_j}$, where $j$ is the $j$-th year ($\mathbf{Z}_{gy_j}$),<br>$\mathbf{Z}_{gy} = \begin{bmatrix} \mathbf{Z}_{gy_1} & \mathbf{0} & \mathbf{0} & \mathbf{0} \\ \mathbf{0} & \mathbf{Z}_{gy_2} & \mathbf{0} & \mathbf{0} \\ \mathbf{0} & \mathbf{0} & \ddots & \mathbf{0} \\ \mathbf{0} & \mathbf{0} & \mathbf{0} & \mathbf{Z}_{gy_J} \end{bmatrix} \text{ and}$ $\text{var}(\mathbf{u}_{gy}) = \mathbf{G}_{gy}$, where $\mathbf{G}_{gy} = \bigoplus_{j=1}^J \mathbf{K}_j\sigma_{gy}^2$,<br>$\mathbf{K}_j$ is the kinship of all entries tested in the $j$-th year. |

\* The subscript $j$ denotes the $j$ -th year; $J$ is the number of years.
† Only applies to Equation 2

**Table A.3.** The variance-covariance structures for each term in Equation 4 for the GCA2 assessment.

| Term and assumption | Variance-covariance structure* | Remarks |
| --- | --- | --- |
| $\mathbf{u}_y \sim N(\mathbf{0}, \mathbf{G}_y)$ | $\mathbf{G}_y = \sigma_y^2 \mathbf{I}$ | Identity variance structure. |
| $\mathbf{u}_{l(y)} \sim N(\mathbf{0}, \mathbf{G}_l)$ | $\mathbf{G}_l = \bigoplus_{j=1}^J \mathbf{G}_{l(j)}$<br>$\mathbf{G}_{l(j)} = \mathbf{I} \sigma_{l(j)}^2$ | Heterogeneous year-specific variance structure in which the diagonal elements $\sigma_{l(j)}^2, \sigma_{t(j)}^2, \sigma_{gt(j)}^2, \sigma_{gl(j)}^2, \sigma_{ls(j)}^2, \sigma_{lsr(j)}^2, \sigma_{lsrb(j)}^2$ differ for $j$-th year for each $\mathbf{G}_{l(j)}, \mathbf{G}_{t(j)}, \mathbf{G}_{gt(j)}, \mathbf{G}_{gl(j)}, \mathbf{G}_{ls(j)}, \mathbf{G}_{lsr(j)}$, and $\mathbf{G}_{lsrb(j)}$, respectively. |
| $\mathbf{u}_{t(y)} \sim N(\mathbf{0}, \mathbf{G}_t)$ | $\mathbf{G}_t = \bigoplus_{j=1}^J \mathbf{G}_{t(j)}$<br>$\mathbf{G}_{t(j)} = \mathbf{I} \sigma_{t(j)}^2$ | |
| $\mathbf{u}_{gt(y)} \sim N(\mathbf{0}, \mathbf{G}_{gt})$ | $\mathbf{G}_{gt} = \bigoplus_{j=1}^J \mathbf{G}_{gt(j)}$<br>$\mathbf{G}_{gt(j)} = \mathbf{I} \sigma_{gt(j)}^2$ | |
| $\mathbf{u}_{gl(y)} \sim N(\mathbf{0}, \mathbf{G}_{gl})$ | $\mathbf{G}_{gl} = \bigoplus_{j=1}^J \mathbf{G}_{gl(j)}$<br>$\mathbf{G}_{gl(j)} = \mathbf{I} \sigma_{gl(j)}^2$ | |
| $\mathbf{u}_{ls(y)} \sim N(\mathbf{0}, \mathbf{G}_{ls})$ | $\mathbf{G}_{ls} = \bigoplus_{j=1}^J \mathbf{G}_{ls(j)}$<br>$\mathbf{G}_{ls(j)} = \mathbf{I} \sigma_{ls(j)}^2$ | |
| $\mathbf{u}_{lsr(y)} \sim N(\mathbf{0}, \mathbf{G}_{lsr})$ | $\mathbf{G}_{lsr} = \bigoplus_{j=1}^J \mathbf{G}_{lsr(j)}$<br>$\mathbf{G}_{lsr(j)} = \mathbf{I} \sigma_{lsr(j)}^2$ | |
| $\mathbf{u}_{lsrb(y)} \sim N(\mathbf{0}, \mathbf{G}_{lsrb})$ | $\mathbf{G}_{lsrb} = \bigoplus_{j=1}^J \mathbf{G}_{lsrb(j)}$<br>$\mathbf{G}_{lsrb(j)} = \mathbf{I} \sigma_{lsrb(j)}^2$ | |
| $\mathbf{u}_g \sim MVN(\mathbf{0}, \mathbf{K} \sigma_g^2)$ | $\mathbf{K} \sigma_g^2$ | $\mathbf{u}_g = \mathbf{Q} \mathbf{v}$, where $\mathbf{Q}$ is the marker genotypes matrix, $\mathbf{v}$ is the vector of marker effects, $\mathbf{v} \sim N(\mathbf{0}, \mathbf{I} \sigma_g^2)$, where $\sigma_g^2$ is the genomic variance.<br>Based on these assumptions $\mathbf{u}_g \sim MVN(\mathbf{0}, \mathbf{K} \sigma_g^2)$, where $\mathbf{K} = \mathbf{Q} \mathbf{Q}^T$. The $\mathbf{K}$ matrix is the genomic relationship matrix. |
| $\mathbf{u}_{gy} \sim MVN(\mathbf{0}, \mathbf{G}_{gy})$ | $\mathbf{G}_{gy} = \bigoplus_{j=1}^2 \mathbf{K}_j \sigma_{gy}^2$ | The matrix $\mathbf{Z}_{gy}$ is a block-diagonal matrix with blocks given by the coefficient of entries in a given year $\bigoplus_{j=1}^2 \mathbf{Z}_{gyj}$,<br>$\mathbf{Z}_{gy} = \begin{bmatrix} \mathbf{Z}_{gy1} & \mathbf{0} \\ \mathbf{0} & \mathbf{Z}_{gy2} \end{bmatrix}$ and $\text{var}(\mathbf{u}_{gy}) = \mathbf{G}_{gy}$,<br>where $\mathbf{G}_{gy} = \bigoplus_{j=1}^2 \mathbf{K}_j \sigma_{gy}^2$,<br>$\mathbf{K}_j$ is the kinship of all entries tested in the $j$-th year. |
| $\mathbf{e} \sim N(\mathbf{0}, \mathbf{R})$ | $\mathbf{R} = \bigoplus_{j=1}^J \mathbf{R}_{l(j)}$<br>$\mathbf{R}_{l(j)} = \mathbf{I} \sigma_{\varepsilon(j)}^2$ | Heterogeneous year×location-specific variance structure consisting of a block-diagonal matrix with diagonal elements $\sigma_{\varepsilon(j)}^2$, $j = 1, 2, \dots, J; l = 1, 2, \dots, L$. |

\* The subscript $j$ denotes the $j$ -th year and subscript $l$ denotes the $l$ -th location. The $J$ is the number of years and $L$ is the number of locations.

## Supplementary Tables

**Table S.1.**
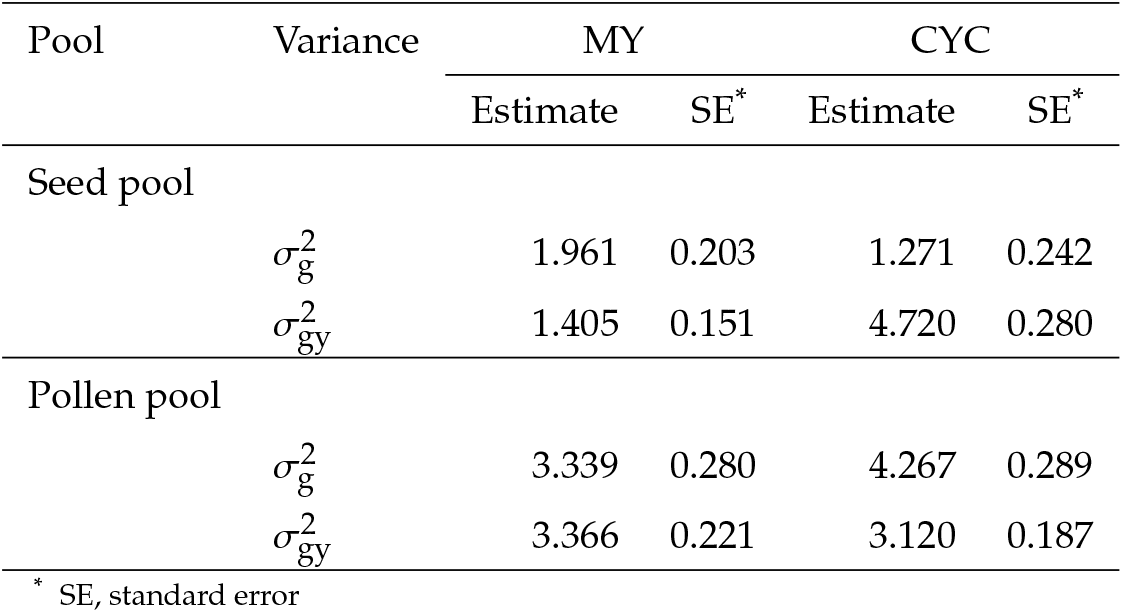
Variance estimates and standard errors for year 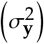, GBV 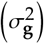, GBV×year interaction effects 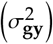 in CYC and MY based on the main year fixed effect model.

**Table S.2.** The 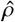 of the CYC and the MY analyses based on the main year fixed effect model.

| Pool | MY |  | CYC |  |
| --- | --- | --- | --- | --- |
| | $\hat{\rho}$ | SE* | $\hat{\rho}$ | SE* |
| Seed pool | 0.582 | 0.044 | 0.212 | 0.038 |
| Pollen pool | 0.498 | 0.032 | 0.578 | 0.026 |

\* SE, standard error. The SE obtained via the Delta method.

**Table S.3.** Variance estimates and standard errors for GBV 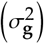 and GBV×year interaction effects 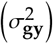 for each cycle of the GCA2 assessment based on the main year fixed effect model.

| Pool | Variance | Cycles |  |  |  |  |  |  |  |
| --- | --- | --- | --- | --- | --- | --- | --- | --- | --- |
|  |  | Cycle 1 |  | Cycle 2 |  | Cycle 3 |  | Cycle 4 |  |
|  |  | Estimate | SE* | Estimate | SE* | Estimate | SE* | Estimate | SE* |
| Seed pool |  |  |  |  |  |  |  |  |  |
| | $\sigma_g^2$ | 0.633 | 0.321 | 0.697 | 0.228 | 1.306 | 0.338 | 0.320 | 0.212 |
| | $\sigma_{gy}^2$ | 0.439 | 0.251 | 0.086 | 0.125 | 0.000 | - | 0.245 | 0.187 |
| Pollen pool |  |  |  |  |  |  |  |  |  |
| | $\sigma_g^2$ | 4.267 | 0.754 | 0.707 | 0.206 | 1.708 | 0.388 | 0.395 | 0.165 |
| | $\sigma_{gy}^2$ | 0.782 | 0.305 | 0.136 | 0.111 | 0.452 | 0.200 | 0.193 | 0.141 |

\* SE, standard error.

